# Humans incorporate attention-dependent uncertainty into perceptual decisions and confidence

**DOI:** 10.1101/175075

**Authors:** Rachel N. Denison, William T. Adler, Marisa Carrasco, Wei Ji Ma

## Abstract

Perceptual decisions are better when they take uncertainty into account. Uncertainty arises not only from the properties of sensory input but also from cognitive sources, such as different levels of attention. However, it is unknown whether humans appropriately adjust for such cognitive sources of uncertainty during perceptual decision making. Here we show that human categorization and confidence decisions take into account uncertainty related to attention. We manipulated uncertainty in an orientation categorization task from trial to trial using only an attentional cue. The categorization task was designed to disambiguate decision rules that did or did not depend on attention. Using formal model comparison to evaluate decision behavior, we found that category and confidence decision boundaries shifted as a function of attention in an approximately Bayesian fashion. This means that the observer’s attentional state on each trial contributed probabilistically to the decision computation. This responsiveness of an observer’s decisions to attention-dependent uncertainty should improve perceptual decisions in natural vision, in which attention is unevenly distributed across a scene.

---

Sensory representations are inherently noisy. In vision, stimulus factors such as low contrast, blur, and visual noise can increase an observer’s uncertainty about a visual stimulus. Optimal perceptual decision-making requires taking into account both the sensory measurements and their associated uncertainty^1^. When driving on a foggy day, for example, you may be more uncertain about the distance between your car and the car in front of you than you would be on a clear day, and try to keep further back. Humans often respond to sensory uncertainty in this way^2,3^, adjusting their choice^4^ behavior as well as their confidence^5^. Confidence is a metacognitive measure that reflects the observer’s degree of certainty about a perceptual decision^6,7^.

Uncertainty arises not only from the external world but also from one’s internal state. Attention is a key internal state variable that governs the uncertainty of visual representations^8,9^; it modulates basic perceptual properties like contrast sensitivity^10,11^ and spatial resolution^12^. Surprisingly, it has been suggested that, unlike for external sources of uncertainty, people fail to take attention into account during perceptual decision-making^13–15^, leading to inaccurate decisions and overconfidence—a risk in attentionally demanding situations like driving a car.

However, this proposal has never been tested using a perceptual task designed to distinguish fixed from flexible decision rules, nor has it been subjected to formal model comparison. Critically, as we show in the **Supplementary Text**, the standard signal detection tasks used previously cannot, in principle, test the fixed decision rule proposal. In standard tasks, the absolute internal decision rule cannot be uniquely recovered, making it impossible to distinguish between fixed and flexible decision rules (**Figure S1a**).

Testing whether observers take attention-dependent uncertainty into account for both choice and confidence also requires a task in which such decision flexibility stands to improve categorization performance. This condition is not met by traditional left versus right categorization tasks, in which the optimal choice boundary is the same (halfway between the means of the left and right category distributions) regardless of the level of uncertainty (**Figure S1b**). Optimal performance can be achieved simply by taking the difference between the evidence for left and the evidence for right, with no need to take uncertainty into account. The same principle applies to present versus absent detection tasks.

To overcome these limitations, we used a recently developed categorization task^4,5^, which we call the *embedded category task*, specifically designed to test whether decision rules depend on uncertainty. In this task, the optimal choice boundaries shift as uncertainty increases, which allowed us to determine whether observers’ behavior tracked these shifts, along with analogous shifts in confidence boundaries. We combined psychophysical experiments with modeling and found that, during perceptual categorization, people not only take attention-dependent uncertainty into account but do so in an approximately Bayesian manner.

## Results

Observers performed the embedded category task, in which they categorized drifting grating stimuli as drawn from either a narrow distribution around horizontal (SD = 3°, category 1) or a wide distribution around horizontal (SD = 12°, category 2) (**Figure 1a**)^4^. This task requires distinguishing a more specific from a more general perceptual category, which is typical of object identification, e.g., distinguishing a beagle from other dogs, an Eames chair from other chairs, or whiskey from other liquids, based on appearance alone. Because the category distributions overlap, maximum accuracy on the task is ~80%.

**Figure 1:**
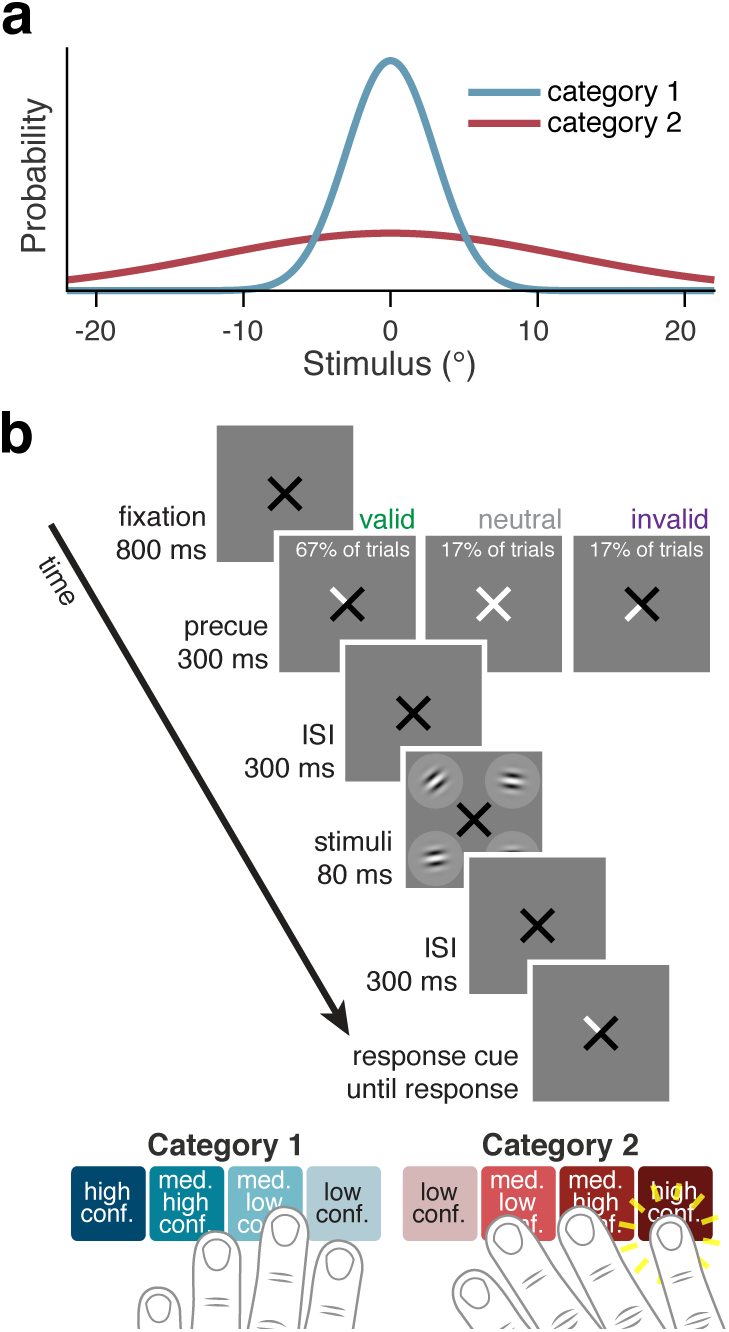
Stimuli and task. (**a**) Stimulus orientation distributions for each category. (**b**) Trial sequence. Cue validity, the likelihood that a precue to one quadrant would match the response cue, was 80%.

We trained observers on the category distributions in category training trials, in which a single stimulus was presented at the fovea, before the main experiment and in short, top-up blocks interleaved with the test blocks (see **Methods**). Accuracy on category training trials in test sessions was 71.9% ± 4.0%, indicating that observers knew the category distributions and could perform the task well.

Four stimuli were briefly presented on each trial, and a response cue indicated which stimulus to report. Observers reported both their category choice (category 1 vs. 2) and their degree of confidence on a 4-point scale using one of 8 buttons, ranging from high-confidence category 1 to high-confidence category 2 (**Figure 1b**). Using a single button press for choice and confidence prevented post-choice influences on the confidence judgment^16^ and emphasized that confidence should reflect the observer’s perception rather than a preceding motor response. We manipulated voluntary (i.e., endogenous) attention on a trial-to-trial basis using a spatial cue that pointed to either one stimulus location (*valid* condition: the response cue matched the cue, 66.7% of trials; and *invalid* condition: it did not match, 16.7% of trials) or all four locations (*neutral* condition: 16.7% of trials) (**Figure 1b**). Twelve observers participated, with about 2000 trials per observer.

Cue validity increased categorization accuracy [one-way repeated-measures ANOVA, *F* (2, 11) = 95.88, *p* < 10^−10^], with higher accuracy following valid cues [two-tailed paired *t*-test, *t*(11) = 7.92, *p* < 10^−5^] and lower accuracy following invalid cues [*t*(11) = 4.62, *p* < 10^−3^], relative to neutral cues (**Figure 2a**, left). This pattern confirms that attention increased orientation sensitivity (e.g.,^11,17^). Attention also increased confidence ratings [*F* (2, 11) = 13.35, *p* < 10^−3^] and decreased reaction time [*F* (2, 11) = 28.76, *p* < 10^−6^], ruling out speed-accuracy tradeoffs as underlying the effect of attention on accuracy (**Figure 2a**).

**Figure 2:**
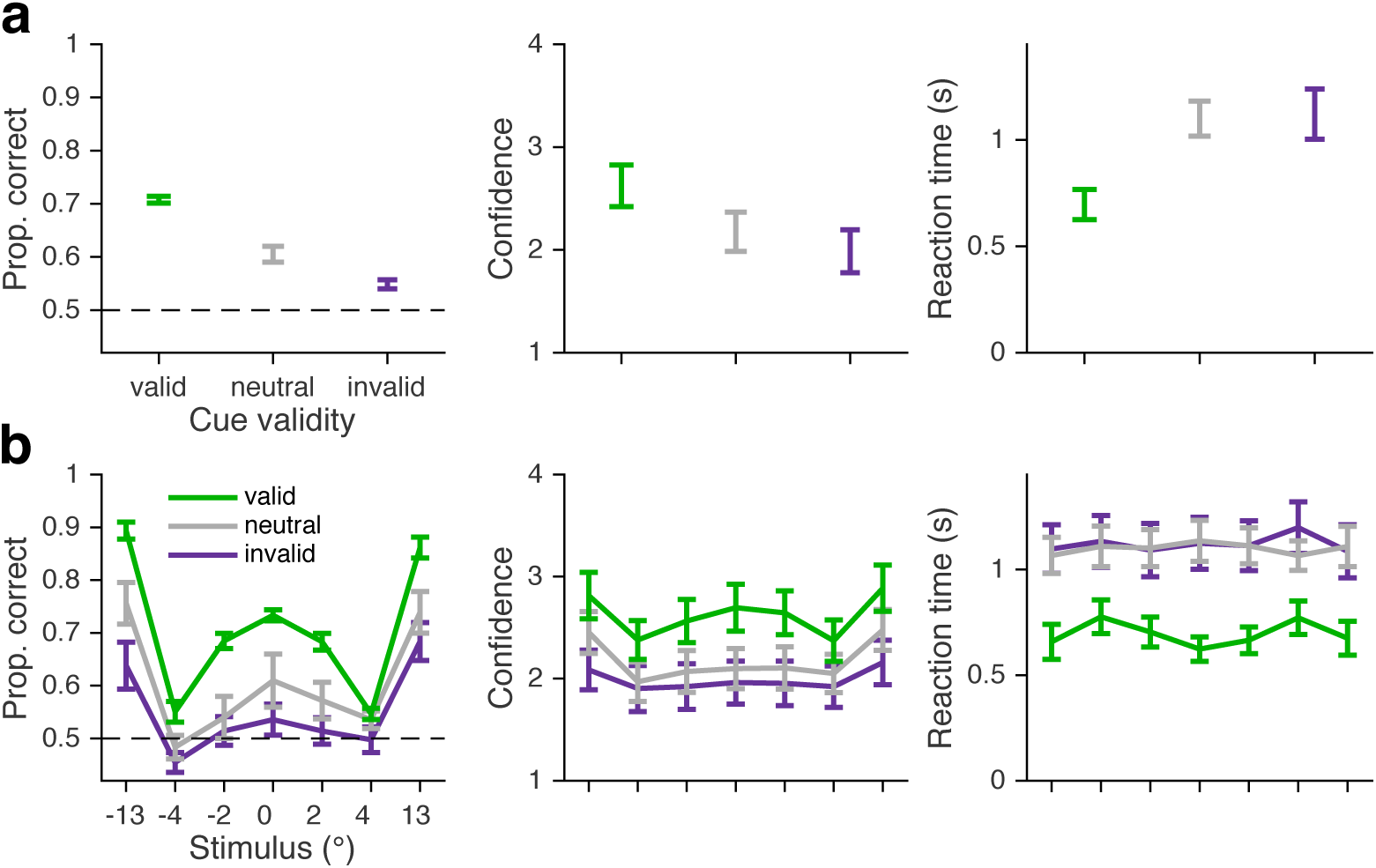
Behavioral data. *n* = 12 observers. Error bars show trial-weighted mean and SEM across observers. (**a**) Accuracy, confidence ratings, and reaction time as a function of cue validity. Maximum accuracy is ~80% because the stimulus distributions overlap. (**b**) As in **a**, but as a function of cue validity and stimulus orientation. Stimulus orientation is binned to equate approximately the number of trials per bin. **Figure S5** shows proportion category 1 choice data and **Figure S6** shows confidence and reaction time data in more detail.

Decision rules in this task are defined by how they map stimulus orientation and attention condition onto a response. We therefore plotted behavior as a function of these two variables. Overall performance was a “W”-shaped function of stimulus orientation (**Figure 2b**, left), reflecting the greater difficulty in categorizing a stimulus when its orientation was near the optimal category boundaries (which were at about 5°). Attention increased the sensitivity of category and confidence responses to the stimulus orientation (**Figure 2b**).

To assess whether observers changed their category and confidence decision boundaries to account for attention-dependent orientation uncertainty, we fit two main models. In one, the Bayesian model, decisions take uncertainty into account, whereas in the other, the Fixed model, decisions are insensitive to uncertainty. Both models assume that, for the stimulus of interest, the observer draws a noisy orientation measurement from a normal distribution centered on the true stimulus value with SD (i.e., uncertainty) dependent on attention. In the Bayesian model, decisions depend on the relative posterior probabilities of the two categories, leading the observer to shift their decision boundaries in measurement space, based on the attention condition^4,5^ (**Figures 3a,b**, **S2**). The Bayesian model maximizes accuracy and produces confidence reports that are a function of the posterior probability of being correct. Note that observers could take uncertainty into account in other ways, but here we began with a normative approach by using a Bayesian model. In the Fixed model, observers use the same decision criteria, regardless of the attention condition^13,15,18–24^ (i.e., they are fixed in measurement space, **Figure 3a,b**). We used Markov Chain Monte Carlo sampling to fit the models to raw, trial-to-trial category and confidence responses from each observer separately (**Methods**, **Table S1**).

**Figure 3:**
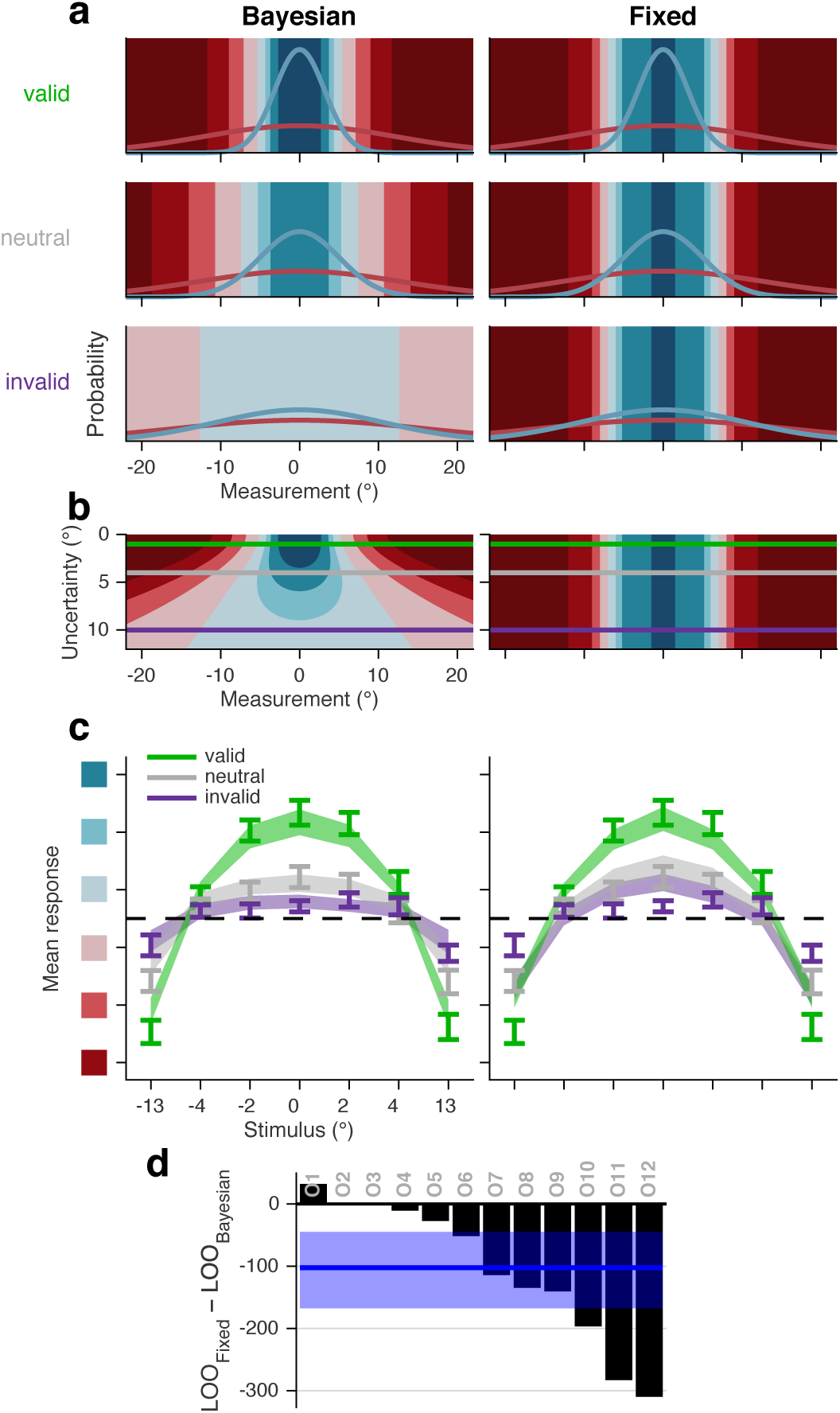
Model schematics, fits, and fit comparison. (**a**) Schematic of Bayesian (left) and Fixed (right) models, which were fit separately for each observer. As attention decreases, uncertainty (the measurement noise SD) increases, and orientation measurement likelihoods (blue and red curves) widen^25^. In the Bayesian model, choice and confidence boundaries are defined by posterior probability ratios and therefore change as a specific function of uncertainty. In the Fixed model, boundaries do not depend on uncertainty. Colors indicate category and confidence response (color code in **Figure 1b**). (**b**) Decision rules for Bayesian and Fixed models show the mappings from orientation measurement and uncertainty to category and confidence responses. Horizontal lines indicate the uncertainty levels used in **a**; note that the regions intersecting with a horizontal line match the regions in the corresponding plot in **a**. (**c**) Model fits to response as a function of orientation and cue validity. Mean response is an 8-point scale ranging from “high confidence” category 1 to “high confidence” category 2, with colors corresponding to those in **Figure 1b**; only the middle 6 responses are shown. Error bars show mean and SEM across observers. Shaded regions are mean and SEM of model fits (**Methods**). Although mean response is shown here, models were fit to raw trial-to-trial data. Stimulus orientation is binned to equate approximately the number of trials per bin. (**d**) Model comparison. Black bars represent individual observer LOO differences of Bayesian from Fixed. Negative values indicate that Bayesian had a higher (better) LOO score than Fixed. Blue line and shaded region show median and 95% confidence interval of bootstrapped mean differences across observers.

Observers’ decisions took attention-dependent uncertainty into account. The Bayesian model captured the data well (**Figure 3c**) and substantially outperformed the Fixed model (**Figure 3c,d**), which had systematic deviations from the data. Although the fit depended on the full data set, note deviations of the Fixed fit from the data near zero tilt and at large tilts in **Figure 3c**, including failure to reproduce the cross-over pattern of the three attention condition curves that is present in the data and the Bayesian fit. To compare models, we used an approximation of leave-one-out cross-validated log likelihood called PSIS-LOO (henceforth LOO)^26^. Bayesian outperformed Fixed by LOO differences (median and 95% CI of bootstrapped mean differences across observers) of 102 [45, 167]. This implies that the attentional state is available to the decision process and is incorporated into probabilistic representations used to make the decision.

Although our main question was whether observers’ decisions took uncertainty into account, our methods also allowed us to determine whether Bayesian computations were necessary to produce the behavioral data, or whether heuristic strategies of accounting for uncertainty would suffice. We tested two models with heuristic decision rules in which the decision boundaries vary as linear or quadratic functions of uncertainty, approximating the Bayesian boundaries (**Figure S3a**). The Linear and Quadratic models both outperformed the Fixed model (LOO differences of 124 [77, 177] and 129 [65, 198], respectively; **Figure S3b,c**). The best model, quantitatively, was Quadratic, similar to previous findings with contrast-dependent uncertainty^4,5^. **Table S2** shows all pairwise comparisons of the models. Model recovery showed that our models were meaningfully distinguishable (**Figure S4**). Decision rules therefore changed with attention without requiring Bayesian computations.

We next asked whether category decision boundaries—regardless of confidence—shift to account for attention-dependent uncertainty. Perhaps, for example, performance of the Bayesian model was superior not because observers changed their categorization behavior, but because they rated their confidence based on the attention condition, which they knew explicitly. Given the mixed findings on the relation between attention and confidence^27–30^, and the proposal that perceptual decisions do not account for attention^13^, such a finding would not be trivial (see **Discussion**); but it would warrant a different interpretation than if category decision boundaries also depended on attention. We fit the four models to the category choice data only and again rejected the Fixed model (**Figure S5a,b**; **Tables S3**, **S4**). Therefore, category criteria, independent of confidence criteria, varied as a function of attention-dependent uncertainty.

Finally, we directly tested for decision boundary shifts—the key difference between the Bayesian and Fixed models—by estimating each observer’s category decision boundaries non-parametrically. To do so, we fit the category choice data with a Free model in which the category decision boundaries varied freely and independently for each attention condition. The estimated boundaries differed between valid and invalid trials (**Figures 4**, **S5c**), with a mean difference of 7.5° (SD = 7.8°) [two-tailed paired *t*-test, *t*(11) = 3.33, *p* < 10^−2^]. Most observers showed a systematic outward shift of category decision boundaries from valid to neutral to invalid conditions, confirming that their choices accounted for uncertainty.

**Figure 4:**
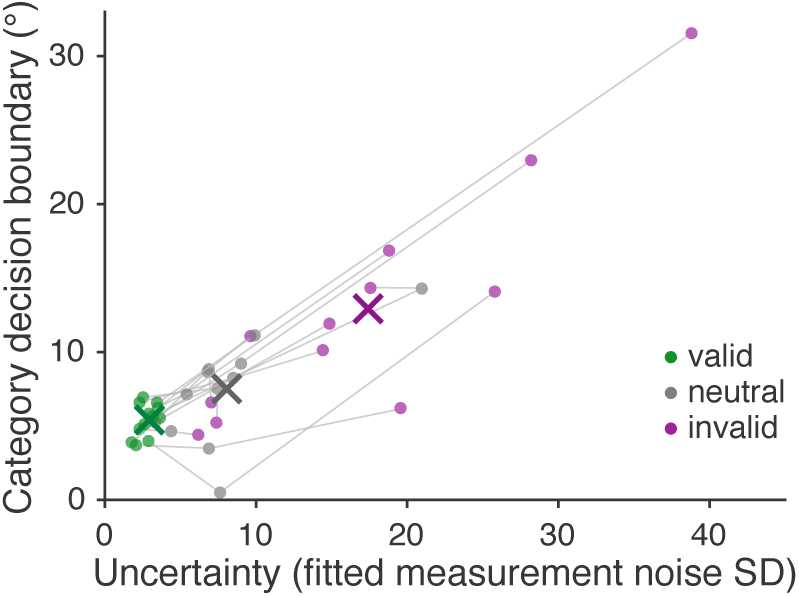
Free model analysis. Group mean MCMC parameter estimates (crosses) show systematic changes in the category decision boundary across attention conditions. The same pattern can be seen for individual observers: each gray line corresponds to a different observer, with connected points representing the estimates for valid, neutral, and invalid attention conditions. Each point represents a pair of parameter estimates: uncertainty and category decision boundary for a specific attention condition.

## Discussion

Using an embedded category task designed to distinguish fixed from flexible decision rules, we found that human perceptual decision-making can take into account uncertainty due to spatial attention. These findings indicate flexible decision behavior that is responsive to attention—an internal factor that affects the uncertainty of stimulus representations.

Our findings of flexible decision boundaries run counter to a previous proposal that observers use a fixed decision rule under varying attentional conditions^13–15,18^. This idea originated from a more general “unified criterion” proposal^22,23^, which asserts that in a display with multiple stimuli, observers adopt a single, fixed decision boundary (the “unified criterion”) for all items^19–24^. The unified criterion proposal implies a rigid, suboptimal mechanism for perceptual decision-making in real-world complex scenes, in which uncertainty can vary due to a variety of factors.

Although the unified criterion proposal has served as a parsimonious explanation for experimental findings^13–15,18–24^, it is impossible to infer decision boundaries from behavior in the signal detection theory (SDT) tasks used previously^31^. In theory, it is always possible to explain behavioral data from such tasks with a fixed decision rule, as long as the means and variances of the internal measurement distributions are free to vary (**Supplementary Text Section S1**).

This issue is particularly thorny for attention studies: SDT works with arbitrary, internal units of “evidence” for one choice or another, and attention could change the means, the variances, or both properties of the internal evidence distributions^10,11,32^. As a result, the decision boundaries are underconstrained: a fixed decision boundary could be mistaken for a flexible one, and vice versa (**Figure 5**). A related point has been made by a study showing that, in a perceptual averaging task, confidence data that appear to be generated by a fixed decision rule can also be explained by a Bayesian decision rule with small underestimations of the internal measurement noise^33^. These considerations underscore the importance of doing model comparison even for relatively simple decision models. It may be, then, that decision boundaries did change with attention in previous studies, but these changes were not inferred for methodological reasons.

**Figure 5:**
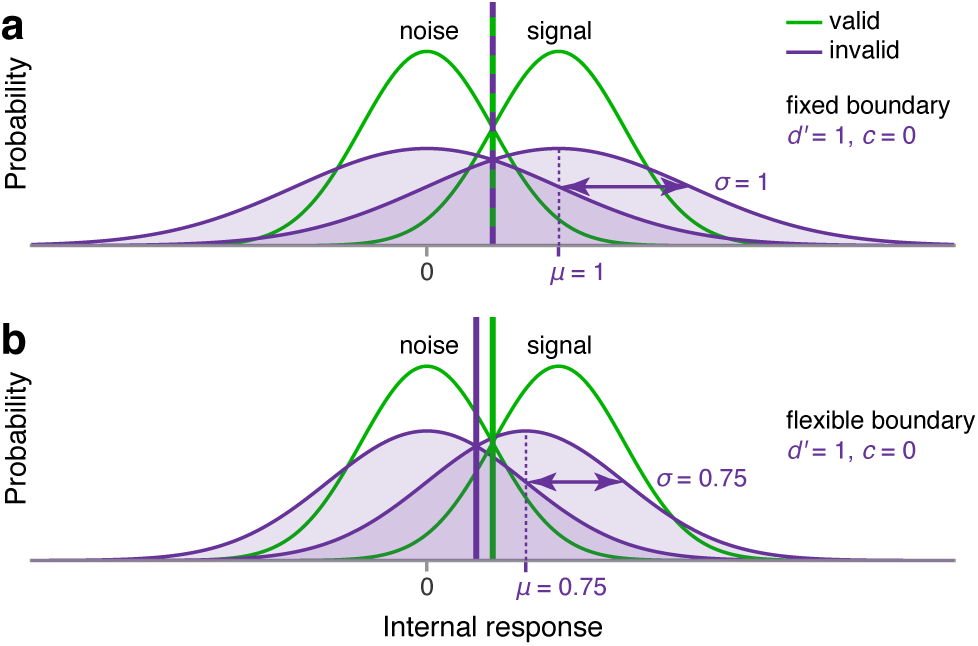
Limitations of standard SDT tasks. SDT tasks such as the detection task illustrated here cannot distinguish fixed from flexible decision rules when the means and variances of internal measurement distributions can also vary across conditions. (**a**) Fixed and (**b**) flexible decision rules give the same behavioral data (perceptual sensitivity, *d*′, and criterion, *c*) in the two depicted scenarios, in which attention affects the measurement distributions differently (compare the invalid distributions in **a** and **b**). An experimenter could not infer from the behavioral data which scenario actually occurred.

Alternatively, it may be that decision boundaries truly did not change in previous studies, and task differences underlie our differing results. Studies supporting the unified criterion proposal employed either detection or orthogonal discrimination^13,15,18–24^, which is often used as a proxy for detection^10,34^. In these tasks, the stimuli are low contrast relative to either a blank screen or a noisy background, and performance is limited by low signal-to-noise ratio. In our categorization task, in contrast, performance is limited by the difficulty of discriminating categories of orientations. Maximum performance depends on the degree of overlap of the category-conditioned stimulus distributions, but variations in performance are determined by the precision of orientation representations, just as in a left vs. right fine discrimination task. Therefore it may be that observers adjust decision boundaries defined with respect to precise features (e.g., what is the exact orientation?) but not boundaries defined with respect to signal strength (e.g., is anything present at all?).

Other task differences could play a role as well. Some previous experiments matched perceptual sensitivity *d*′ for different attention conditions by changing stimulus contrast; in these experiments, attention and physical stimulus properties varied together^13,15^. For the metacognitive report, our study asked for confidence rather than visibility^13^; these subjective measures are known to differ^35^. Finally, one study^15^ using a signal detection approach suggested that observers do not fully take into account an instructed prior, and take it into account less when attention is low. The question of how attention affects the use of a prior is different from the question asked in the current study, as incorporating a prior requires a cognitive step beyond accounting for uncertainty in the perceptual representation. In the future, it will be interesting to examine how decision boundaries relate to explicit priors using tasks in which absolute decision boundaries can be uniquely inferred.

Despite the fact that attention has a large influence on visual perception^8^, only a handful of studies have examined the influence of attention on confidence. Their findings have been mixed. Two studies found that voluntary attention increased confidence^27,28^; one found that voluntary but not involuntary attention increased confidence^30^; and another found no effect of voluntary attention on confidence^29^. This last result has been attributed to response speed pressures^27,30^. Three other studies suggested an inverse relation between attention and confidence, though these used rather different attention manipulations and measures. One study reported higher confidence for uncued compared to cued error trials^36^; one found higher confidence for stimuli with incongruent compared to congruent flankers^37^; and a third found that lower fMRI BOLD activation in the dorsal attention network correlated with higher confidence^18^. Our results, based on an experimental manipulation of spatial attention with no response speed pressure, support a positive relation between spatial attention and confidence and further reveal that it is approximately Bayesian.

The mechanisms for decision adjustment under attention-dependent uncertainty could be mediated by effective contrast^10,38,39^. Alternatively, attention-dependent decision-making may rely on higher-order monitoring of attentional state. For example, the observer could consciously adjust a decision depending on whether he or she was paying attention. Future studies will be required to distinguish between these more bottom-up or top-down mechanisms.

Our finding that human observers incorporate attention-dependent uncertainty into perceptual categorization and confidence reports in a statistically appropriate fashion points to the question of what other kinds of internal states can be incorporated into perceptual decision-making. There is no indication, for example, that direct stimulation of sensory cortical areas leads to adjustments of confidence and visibility reports^18,40,41^, suggesting that the system is not responsive to every change to internal noise. It may be that the system is more responsive to states that are internally generated or that have consistent behavioral relevance. Attention is typically spread unevenly across multiple objects in a visual scene, so the ability to account for attention likely improves perceptual decisions in natural vision. It remains to be seen whether the perceptual decision-making system is responsive to other cognitive or motivational states.

## ACKNOWLEDGEMENTS

The authors would like to thank: Roshni Lulla and Gordon C. Bill for assistance with data collection and helpful discussions; Luigi Acerbi for helpful ideas and tools related to model fitting and model comparison; and Stephanie Badde, Michael Landy, and Christopher Summerfield for comments on the manuscript. This material is based upon work supported by the National Science Foundation Graduate Research Fellowship under Grant No. DGE-1342536 (W.T.A.) and by the National Eye Institute of the National Institutes of Health under awards T32EY007136 and F32EY025533 (R.N.D.).

## AUTHOR CONTRIBUTIONS

R.N.D, W.T.A., M.C., and W.J.M. designed the research and wrote the manuscript. W.T.A. and W.J.M. developed the models. R.N.D. and W.T.A. performed the experiments and analyzed data.

## COMPETING FINANCIAL INTERESTS

The authors declare no competing financial interests.

## DATA AND CODE AVAILABILITY

All data and code used for running experiments, model fitting, and plotting is available on GitHub (https://github.com/wtadler/confidence).

## Methods

### 1 Experiment

#### 1.1 Observers

Twelve observers (7 female, 5 male), aged 18–25 years, participated in the experiment. These observers came from an original set of 28 observers who completed at least one session. The remaining observers did not complete the main experiment, either because they were not invited to continue following the pre-screening staircase sessions (15 observers, **Section 1.3.7**) or because they chose to stop participating before all sessions were completed (one observer). Observers received $10 per 40–60 minute session, plus a completion bonus of $25. The experiments were approved by the University Committee on Activities Involving Human Subjects of New York University. Informed consent was given by each observer before the experiment. All observers were naïve to the purpose of the experiment. No observers were fellow scientists.

#### 1.2 Apparatus and stimuli

##### 1.2.1 Apparatus

Observers were seated in a dark room, at a viewing distance of 57 cm from the screen, with their chin in a chinrest. Stimuli were presented on a gamma-corrected 100 Hz, 21-inch display (Model Sony GDM-5402). The display was connected to a 2010 iMac running OS X 10.6.8 using MATLAB (Mathworks) with Psychophysics Toolbox 3^42–44^.

##### 1.2.2 Stimuli

The background was mid-level gray (60 cd/m^2^). Stimuli consisted of drifting Gabors with a spatial frequency of 0.8 cycles per degree, a speed of 6 cycles/s, a Gaussian envelope with a SD of 0.8 degrees of visual angle (dva), and a randomized starting phase. In category training, the stimuli were positioned at fixation, and the central fixation cross was a black “+” subtending 1.2 dva in diameter. In all other blocks, one stimulus was positioned in each of the four quadrants of the screen, at 45, 135, 225, and 315 degrees, 5 dva from fixation, and the fixation cross was a black “×” with each arm pointing to a quadrant. One or more of the arms turned white to provide a precue or response cue (**Figure 1b**). Stimulus contrast depended on the block type.

##### 1.2.3 Categories

Stimulus orientations *s_i_* were drawn from Gaussian distributions with means *µ*_1_ = *µ*_2_ = 0°, and standard deviations *σ*_1_ = 3° (category 1) and *σ*_2_ = 12° (category 2). Because the category distributions overlapped, maximum accuracy was ~80%.

##### 1.2.4 Attention manipulation

During attention training and testing blocks, voluntary spatial attention was manipulated via a central precue presented at the start of the trial. A response cue at the end of the trial indicated which of the four stimuli to report. On each trial, each of the four stimuli was drawn from one of the two category distributions. Each stimulus was generated independently. In valid trials (66.7% of all trials), a single quadrant was precued and the response cue matched the precue. In invalid trials (16.7%), a single quadrant was precued and the response cue did not match the precue. Cue validity was therefore 80% when a single quadrant was precued. In neutral trials (16.7%), all four quadrants were precued, and the response cue pointed to one of the four quadrants with equal probability for each quadrant.

#### 1.3 Procedure

Each observer completed seven sessions. Because our behavioral task involved multiple components—orientation categorization, confidence reports, and attention—we trained observers on each component in a stepwise fashion, as described below.

The first two sessions (“staircase sessions”) were used to pre-screen observers and find a stimulus contrast level that would achieve maximum separability in performance across the three attention conditions. Each staircase session consisted of 3 category training blocks and 3 category/attention testing-with-staircase blocks, in alternation. No confidence reports were collected in these sessions. The first category training block was preceded by a category demo, and the first category/attention testing-with-staircase block was preceded by a category/attention training block. Detailed instructions were provided in the first session. Most blocks consisted of sets of trials, in between which the observer was informed of their progress (e.g., “You have completed three quarters of Testing Block 2 of 3”) and allowed to rest. The staircase sessions also served as practice on the categorization and attention components of the task, so that observers knew them well by the time they started the main experiment. During these sessions, stimulus contrast was 35% for training blocks, and varied during the testing-with-staircase blocks.

The final five sessions (“test sessions”) comprised the main experiment. Each test session consisted of 3 category training blocks and 3 confidence/attention testing blocks, in alternation. The first category training block was preceded by a category demo, and the first confidence/attention testing block was preceded by a confidence/attention training block. During these sessions, stimulus contrast was fixed to an observer-specific value in all blocks.

Combining all test sessions, 9 observers completed 15 confidence/attention testing blocks (2160 trials), 2 observers completed 14 testing blocks (2016 trials), and 1 observer completed 12 testing blocks (1728 trials). Accuracy on category training trials was 70.8% ± 4.0% (mean ± 1 SD) in staircase sessions and 71.9% ± 4.0% in test sessions, indicating that observers learned the category distributions well (recall that maximum accuracy on the task is ~80%).

##### 1.3.1 Eye tracking

Eye tracking (Eyelink 1000) was used to monitor fixation online. In all blocks, trials were only initiated when the observer was fixating. In testing blocks, trials in which observers broke fixation due to blinks or eye movements were aborted and repeated later in the experiment.

##### 1.3.2 Instructions

*First staircase session*. Before the first category training block, we provided observers with a printed graphic similar to **Figure 1a**, explained how the stimuli were generated from distributions, and explained the category training procedure. We also explained that trials would only proceed when the observer maintained fixation. Before the category/attention training block, we explained the attention task using an onscreen graphic that explained the cuing procedure and a printed graphic that illustrated cue validity. We also explained the requirement to maintain fixation from the precue until the response cue and the consequences of breaking fixation. Before the first category/attention testing-with-staircase block, we explained that the stimulus presentation time would be shorter and that the contrast of the stimuli would vary.

*First test session*. Before the confidence/attention training block, we explained two changes to the experiment. First, we told observers that they would be reporting category choice and confidence simultaneously. We provided a printed graphic similar to the buttons shown in **Figure 1b**, showing the eight buttons representing category choice and confidence level, the latter on a 4-point scale. The confidence levels were labeled as “very high,” “somewhat high,” “somewhat low,” and “very low.” All printed graphics were visible to observers throughout the experiment. Second, we told observers that contrast would be fixed (rather than variable) for the remainder of the experiment, in all blocks.

##### 1.3.3 Category demo

We showed observers 25 randomly drawn exemplar stimuli from each category (50 exemplars in the first staircase session). Stimulus contrast was 35% in staircase sessions and observer-specific in test sessions.

##### 1.3.4 Category training

To ensure that observers knew the stimulus distributions well, we gave them extensive category training with trial-to-trial correctness feedback and foveal stimulus presentation to reduce orientation uncertainty. Each trial proceeded as follows: Observers fixated on a central cross for 1 s. Category 1 or category 2 was selected with equal probability. The stimulus orientation was drawn from the corresponding stimulus distribution and displayed as a drifting Gabor. The stimulus appeared at fixation for 300 ms, replacing the fixation cross. Observers were asked to report category 1 or category 2 by pressing a button with their left or right index finger, respectively. Observers were able to respond immediately after the offset of the stimulus, at which point correctness feedback was displayed for 1.1 s, e.g., “You said Category 1. Correct!” The fixation cross then reappeared. In staircase sessions, the stimulus contrast was 35%. In test sessions, the contrast matched the observer-specific levels chosen for testing blocks, in order to minimize obvious changes between training and testing blocks. Each category training block had 2 sets of 36 trials (72 total). At the end of the block, observers were shown the percentage of trials that they had correctly categorized.

##### 1.3.5 Category/attention training

To familiarize observers with the attention task before the testing-with-staircase blocks, they completed category/attention training. Observers performed the attention task, reporting only category choice. To prevent observers from forming a simple mapping of orientation measurement and attention condition onto the probability of category 1 (which might have biased behavior towards the Bayesian model), we withheld trial-to-trial feedback on this and all other types of attention blocks. The precue indicating which location(s) to attend to appeared for 300 ms, followed by a 300 ms period in which a standard fixation cross was shown. Then the four drifting Gabor stimuli were displayed for 300 ms. After another 300 ms period with a fixation cross, the response cue appeared, indicating which stimulus to report. The response cue remained on the screen until the observer pressed one of the two choice response buttons, with no time pressure. Observers were free to blink or rest briefly between trials, with a minimum intertrial interval of 800 ms. All attention conditions were randomly intermixed. The stimulus contrast was 35%, as in staircase session category training. The block had 36 trials in the first session and 30 trials in subsequent sessions. At the end of the block, observers were shown the percentage of trials they had correctly categorized.

##### 1.3.6 Category/attention testing-with-staircase

The purpose of this block was to determine the stimulus contrast for each observer that would be used in the test sessions. The trial procedure was identical to that of category/attention training, except that stimulus presentation time was 80 ms (instead of 300 ms) and stimulus contrast varied. We used an adaptive staircase procedure to determine the stimulus contrast on each trial and estimate psychometric functions for performance accuracy as a function of log contrast. Separate staircases were used for valid, neutral, and invalid conditions. We used Luigi Acerbi’s MATLAB (https://github.com/lacerbi/psybayes) implementation of the PSI method by Kontsevich and Tyler^45^, extended to include the lapse rate^46^. The method generates a posterior distribution over three parameters of the psychometric function: threshold *µ*, slope *σ*, and lapse rate *λ*. On each trial, it selects a stimulus intensity that maximizes the expected information gain by completion of the trial. *µ* (log contrast units) ranged from −6.5 to 0 and had a Gaussian prior distribution with mean −2 and SD 1.2. log *σ* ranged from −3 to 0, and had a uniform prior distribution across the range. *λ* ranged from 0.15 (because the maximum accuracy in the task was slightly below 1 − 0.15) to 0.5, and had a Beta prior distribution with shape parameters *α* = 20 and *β* = 39. Each block had 4 sets of 36 trials (144 total). At the end of the block, observers were shown the percentage of trials that they had correctly categorized.

##### 1.3.7 Observer pre-screening and contrast selection

Simulations we conducted before starting the study showed that without a sufficiently large noise (related to accuracy) difference between valid and invalid trials, our models would be indistinguishable. Therefore, we used a pre-screening process to select observers with a robust attention effect to participate in the main experiment. We also determined the stimulus contrast at which each observer’s attention effect was maximal. This procedure increased the probability that uncertainty would depend on attention in the main experiment, which was critical for answering our central question about decision behavior. Note that the pre-screening procedure only concerned the overall accuracy difference between valid and invalid trials, which is independent of how attention affects the decision rule.

After each observer’s final staircase session, we plotted and visually inspected the mean and SD of the posterior over the 3 (valid, neutral, and invalid) estimated psychometric functions (an example is shown in **Figure S7**). An observer was considered eligible for the remainder of the study if there existed a contrast that satisfied two conditions. 1) Invalid accuracy was above chance: The mean minus the SD of the posterior over invalid psychometric functions was above 0.5. 2) Valid accuracy was different from invalid accuracy: The mean minus the SD of the posterior over valid psychometric functions was greater than the mean plus 1 SD of the posterior over invalid psychometric functions. For example, note that there is a range of values in **Figure S7** for which the purple shading does not overlap with the chance line or with the green shading. Within the range of suitable contrasts, we selected the contrast for which the separation between valid, neutral, and invalid performance appeared to be maximal. Observers for which no suitable contrast could be found were not invited to participate in the main experiment. Selected contrasts ranged from 4% to 60% across observers.

##### 1.3.8 Confidence/attention training

To familiarize observers with the button mappings for choice and confidence, they completed confidence/attention training. The trial procedure was identical to category/attention training, except observers reported their confidence on each trial in addition to their category choice. Observers were not instructed to use the full range of confidence reports, as that might have biased them away from reporting what felt most natural. Instead, they were simply asked to be “as accurate as possible in reporting their confidence” on each trial. Feedback about their choice and confidence report was presented for 1.2 s after each trial, e.g. “You said category 2 with HIGH confidence.” The stimulus contrast was specific to each observer, based on the staircase sessions. There were 30 trials per block.

##### 1.3.9 Confidence/attention testing

These were the main experimental blocks. The trial procedure (**Figure 1b**) was the same as in confidence/attention training blocks, but with no trial-to-trial feedback whatsoever. Each block had 4 sets of 36 trials (144 total). At the end of each block, observers were required to take a break of at least 30 s. During the break, they were shown the percentage of trials that they had correctly categorized. Observers were also shown a list of the top 10 block scores (across all observers, indicated by initials). This was intended to motivate observers to perform well, and to reassure them that their scores were normal, since it is rare to score above 75% on a block.

### 2 Modeling

The modeling procedures were similar to those used by Adler and Ma^5^. Several modeling choices were adopted based on model comparisons performed for that study. These included: having orientation-dependent measurement noise; allowing all decision boundaries to be free parameters in the Bayesian model; including decision noise in the Bayesian model; and modeling three types of lapse rates.

#### 2.1 Measurement noise

We used free parameters to characterize *σ*, the standard deviation (SD) of orientation measurement noise, for all three attention conditions: *σ*_valid_, *σ*_neutral_, and *σ*_invalid_.

We assumed additive orientation-dependent noise in the form of a rectified 2-cycle sinusoid, accounting for the finding that measurement noise is higher at noncardinal orientations^47^. For a given trial *i*, the measurement noise SD comes out to

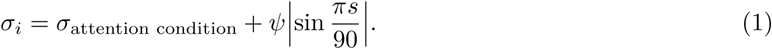

The second term of this equation is a constant that depends on the stimulus orientation *s*, with *ψ* a free parameter that determines the degree of orientation dependence.

#### 2.2 Response probability

We coded all responses as *r* ∈ {1, 2, …, 8}, with each value indicating category and confidence. A value of 1 mapped to high confidence category 1, and a value of 8 mapped to high confidence category 2, as in **Figure 1b**. The probability of a single trial *i* is equal to the probability mass of the internal measurement distribution 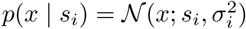 in a range corresponding to the observer’s response *r_i_*. Because we only use a small range of orientations, we can safely approximate measurement noise as a normal distribution, rather than a von Mises distribution. We find the boundaries 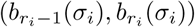 in measurement space, as defined by the fitting model *m* and parameters *θ*, and then compute the probability mass of the measurement distribution between the boundaries:

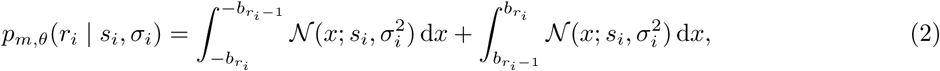

where *b*_0_ = 0° and *b*_8_ = ∞°.

To obtain the log likelihood of the dataset, given a model with parameters *θ*, we compute the sum of the log probability for every trial *i*, where *t* is the total number of trials:

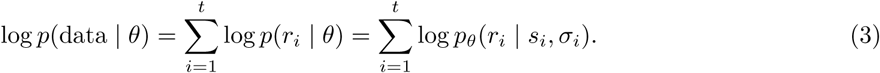

#### 2.3 Model specification

##### 2.3.1 Bayesian

*Derivation of d*. The log posterior ratio *d* is equivalent to the log likelihood ratio plus an additive term representing the prior probability over category:

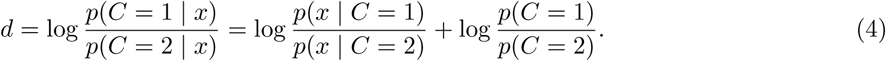

To get *d*, we need to find the expressions for the orientation measurement likelihood *p*(*x* | *C*). The observer knows that the measurement *x* is caused by the stimulus *s*, but has no knowledge of *s*. Therefore, the optimal observer marginalizes over *s*:

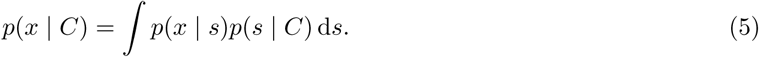

We substitute the expressions for the noise distribution and the stimulus distribution, and evaluate the integral:

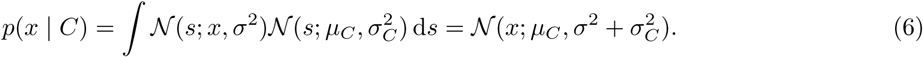

Plugging in the category-specific *µ_C_* and *σ_C_*, and substituting these expressions back into equation (4), we get:

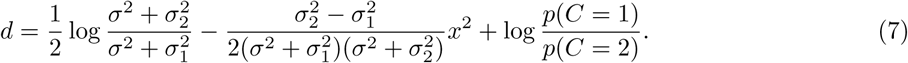

The 8 possible category and confidence responses are determined by comparing the log posterior ratio *d* to a set of decision boundaries **k** = (*k*_0_, *k*_1_, …, *k*_8_). *k*_4_ is equal to the observer’s believed log prior ratio 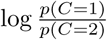, which functions as the boundary on *d* between the 4 category 1 responses and the 4 category 2 responses and is fit to capture possible category bias. *k*_4_ is the only boundary parameter in models of category choice only (and not confidence). *k*_0_ is fixed at −∞ and *k*_8_ is fixed at ∞. The observer chooses category 1 when *d* is positive. Thus there were 7 free boundary parameters: *k*_1_,*k*_2_,*k*_3_,*k*_4_,*k*_5_,*k*_6_,*k*_7_. The posterior probability of category 1 can be written as as 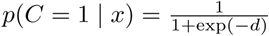.

*Decision boundaries*. In the Bayesian models with *d* noise, we assume that, for each trial, there is an added Gaussian noise term on *d*, *η_d_* ~ *p*(*η_d_*), where 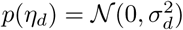, and *σ_d_* is a free parameter. We pre-computed 101 evenly spaced draws of *η_d_* and their corresponding probability densities *p*(*η_d_*). We used equation (7) to compute a lookup table containing the values of *d* as a function of *x*, *σ*, and *η_d_*. We then used linear interpolation to find sets of measurement boundaries **b**(*σ*) corresponding to each draw of *η_d_*^48^. We then computed 101 response probabilities for each trial (as described in **Section 2.2**), one for each draw of *η_d_*, and computed the weighted average according to *p*(*η_d_*). This gave the values of *p_m_*,*_θ_*(*r_i_* | *s_i_*,*σ_i_*) for each trial *i*, which are needed in order to compute the total log likelihood of the dataset under the model.

In the Bayesian choice model without *d* noise, we translate the decision boundary *k*_4_ from a log prior ratio to a measurement boundary corresponding to the fitted noise levels *σ*. To do this, we use *k*_4_ as the left-hand side of equation (7) and solve for *x* at the fitted levels of *σ*. We used this model only for the purpose of obtaining estimates of the category decision boundary parameters, and not for model comparison.

##### 2.3.2 Fixed

In the Fixed model, the observer compares the measurement to a set of boundaries that are not dependent on *σ*. We fit free parameters **k** and use measurement boundaries *b_r_* = *k_r_*.

##### 2.3.3 Linear and Quadratic

In the Linear and Quadratic models, the observer compares the measurement to a set of boundaries that are linear or quadratic functions of *σ*. We fit free parameters **k** and **m** and use measurement boundaries *b_r_*(*σ*) = *k_r_* + *m_r_σ* (Linear) or *b_r_*(*σ*) = *k_r_* + *m_r_σ*^2^ (Quadratic).

##### 2.3.4 Free

To estimate the category boundaries with minimal assumptions, we fit a Free model in which the observer compares the orientation measurement to a set of boundaries that vary nonparametrically (i.e., free of a parametric relationship with *σ*) across attention conditions. As with the Bayesian choice model without *d* noise (**Section 2.3.1**), we used this model only for the purpose of obtaining estimates of the category decision boundary parameters and did not fit confidence. We fit free parameters *k*_4,valid_, *k*_4,neutral_, *k*_4,invalid_, and used measurement boundaries *b*_4,attention condition_ = *k*_4,attention condition_.

#### 2.4 Model fitting

Rather than find a maximum likelihood estimate of the parameters, we sampled from the posterior distribution over parameters, *p*(*θ* | data); this has the advantage of maintaining a measure of uncertainty about the parameters, which can be used both for model comparison and for plotting model fits. To sample from the posterior, we use an expression for the unnormalized log posterior

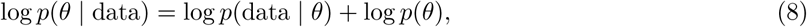

where log *p*(data | *θ*) is given in equation (3). We assumed a factorized prior over each parameter *j*:

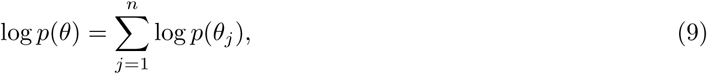

where *j* is the parameter index and *n* is the number of parameters. We took uniform (or, for parameters that were standard deviations, log-uniform) priors over reasonable, sufficiently large ranges^48^, which we chose before fitting any models.

We sampled from the probability distribution using a Markov Chain Monte Carlo (MCMC) method, slice sampling^49^. For each model and dataset combination, we ran between 4 and 10 parallel chains with random starting points. For each chain, we took 100,000 to 1,000,000 total samples (depending on model computational time) from the posterior distribution over parameters. We discarded the first third of the samples and kept 6,667 of the remaining samples, evenly spaced to reduce autocorrelation. All samples with log posteriors more than 40 below the maximum log posterior were discarded. Marginal probability distributions of the sample log likelihoods were visually checked for convergence across chains. In total we had 120 model and dataset combinations, with a median of 40,002 kept samples (interquartile range = 13,334).

#### 2.5 Model comparison

##### 2.5.1 Metric choice

To compare model fits while accounting for the complexity of each model, we computed an approximation of leave-one-out cross-validation. Leave-one-out cross-validation is the most thorough way to cross-validate but is very computationally intensive; it requires fitting the model *t* times, where *t* is the number of trials. The Pareto smoothed importance sampling approximation of leave-one-out cross-validation (PSIS-LOO, referred to here simply as LOO) takes into account the model’s uncertainty landscape by using samples from the full posterior of *θ*^26^. LOO is currently the most accurate approximation of leave-one-out cross-validation^50^.

##### 2.5.2 Metric aggregation

In all figures where we present model comparison results (**Figures 3d**, **S3c**, **S5b**), we aggregate LOO scores by the following procedure: Choose a reference model (e.g. Fixed). Subtract all LOO scores from the corresponding observer’s score for that model; this converts all scores to a LOO “difference from reference” score, with lower (more negative) indicating a better score and higher (more positive) indicating a worse score. Repeat the following standard bootstrap procedure 10,000 times: Choose randomly, with replacement, a group of datasets equal to the total number of unique datasets, and take the mean of their “difference from reference” scores for each model. Blue lines and shaded regions in model comparison plots indicate the median and 95% CI on the distribution of these bootstrapped mean “difference from reference” scores.

## Supplementary Text

### S1 Theoretical motivations for using the embedded category task

The goal of the current study was to test whether category and confidence decision rules account for attention-dependent uncertainty. The embedded category task^4^ can answer this question, whereas standard signal detection theory^51^ (SDT) detection and coarse discrimination (e.g., ±45°) tasks are unable to do so. There are two key theoretical advantages of the embedded category task over the SDT tasks: 1) the ability to infer absolute decision boundaries, and 2) the incentive to shift the category boundary when uncertainty changes.

#### S1.1 Inference of absolute decision boundaries

A decision rule can be thought of as a boundary defined on the observer’s internal measurement space. Here we were interested in the absolute location of that boundary *b*. The “unified criterion” discussed previously also refers to an absolute boundary^22,23^.

The embedded category task can be used to infer absolute decision boundaries from behavioral data, because the measurement axis represents known feature values – specifically, orientation values. Making a category or confidence decision can be thought of as comparing the observed stimulus orientation to an internal reference orientation, which is the decision boundary. As experimenters, we know the means of the internal measurement distributions (specific orientations), so we can infer the absolute decision boundary on the orientation axis.

In SDT detection and coarse discrimination tasks, in contrast, absolute decision boundaries cannot be inferred, because the measurement axis represents values that we, as experimenters, do not know. In a detection task, the measurement value is thought of as the strength of the internal signal, or the “amount of evidence” that the external signal is present. In a coarse discrimination task, the measurement value is thought of as the amount of evidence for choice 1 (e.g., −45°) versus choice 2 (e.g., +45°). We don’t know the means of the internal measurement distributions in real values; we don’t even know what the units are. Consequently, the behavioral SDT measures *d*′ (perceptual sensitivity) and *c* (criterion) are defined in a normalized space – *d*′ and *c* are z-scored measures of the distance between the two internal category distributions and the location of the observer’s decision boundary, respectively. So they are relative measures.

As a result, an absolute decision boundary *b* is unrecoverable from behavioral data. This fact can be shown mathematically. The standard formulae for *d*′ and *c* are

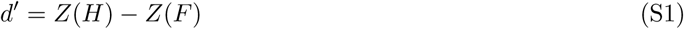

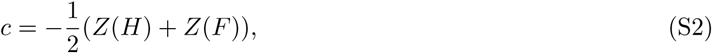

where *Z* is the inverse of the normal cumulative distribution function (i.e., z-score), *H* is the proportion of hits, and *F* is the proportion of false alarms. Note this formula gives *c* with respect to the unbiased criterion. If we let the mean of the noise distribution be 0 and the mean of the signal distribution be *µ*, then

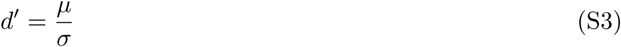

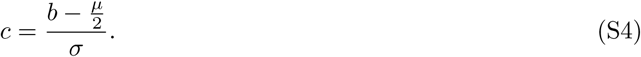

Here we have two equations with three unknowns. Any combination of *d*′ and *c* is therefore consistent with an infinite set of combinations of the *µ*, *σ*, and *b* parameters; thus *b* cannot be uniquely determined. The intuition here is that the SDT axis can be rescaled without changing *d*′ and *c* (**Figure S1a**). The same issue applies not only to *d*′ and *c* but to any other relative behavioral measure, such as hit rate or false alarm rate. Kontsevich et al.^31^ raised this concern about Gorea and Sagi’s^23^ proposal of a unified criterion for simultaneously presented stimuli.

**Figure S1:**
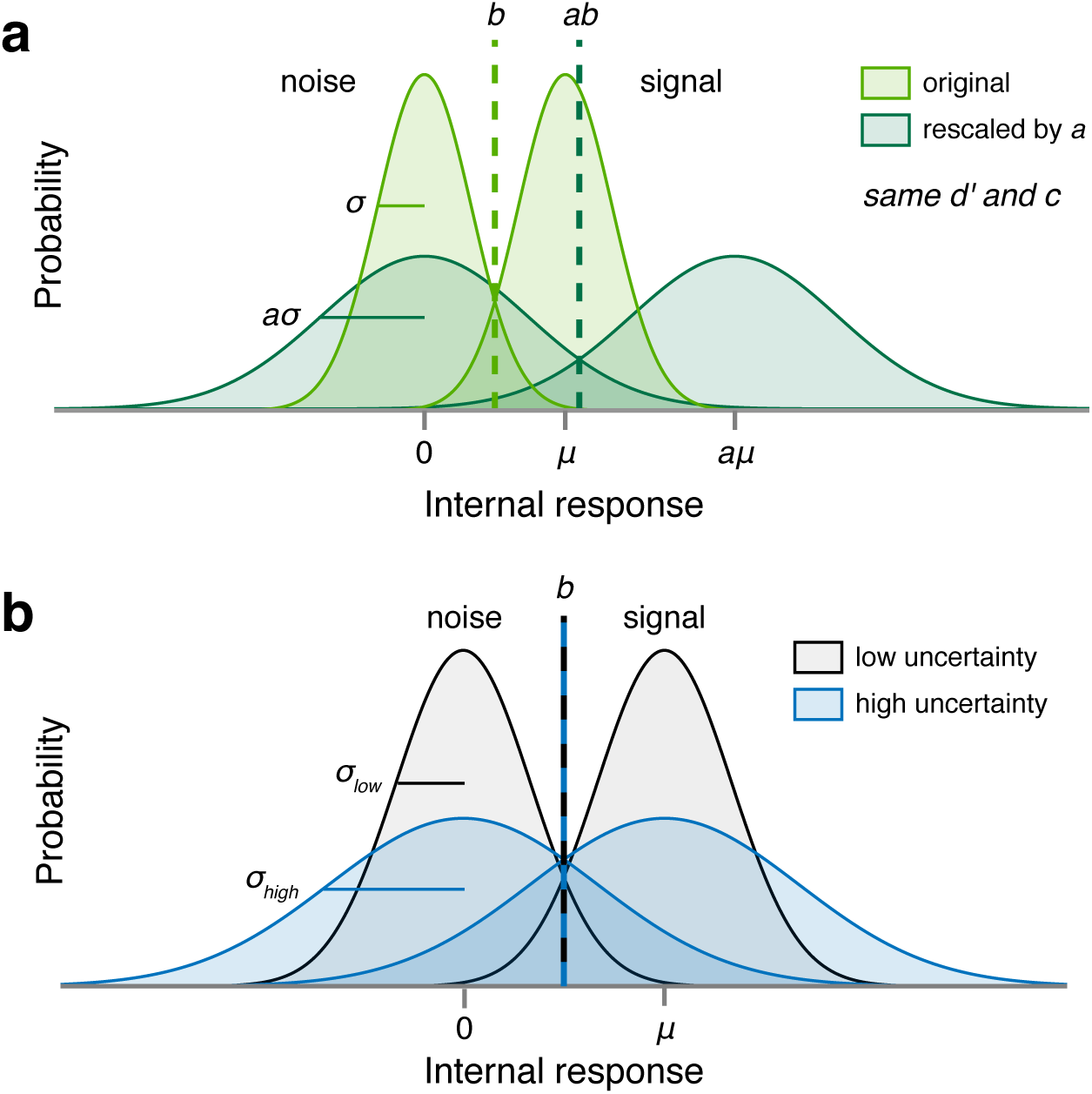
Methodological limitations in standard signal detection tasks. (**a**) Rescaling the SDT axis by a factor *a* yields the same values of *d*′ and *c*, but with a different set of parameters (the original parameters rescaled by *a*). This is because *d*′ and *c* are relative to the internal measurement distributions, not any absolute evidence metric. (**b**) In standard SDT tasks, when the means of the internal measurement distributions are symmetric about the optimal category boundary, changing the uncertainty does not change the optimal boundary. *µ* = mean, *σ* = standard deviation, *b* = decision boundary.

The non-uniqueness of SDT parameters creates a critical problem when asking whether *b* changes with attention. Attention could change *µ*, *σ*, or both properties of the internal measurement distributions^10,11,32^. Therefore, *b* cannot be compared, even in a relative fashion, across attention conditions; so fixed and flexible decision rules cannot be distinguished (**Figure 5**).

Note that changes in *confidence* boundaries with uncertainty can be determined even for a standard SDT task as long as a known stimulus dimension is manipulated simultaneously. For example, a left vs. right fine discrimination task in which stimuli are drawn from orientation distributions with similar means can be used to infer absolute confidence boundaries on an orientation axis^5^. In general, two-dimensional data (e.g., orientation × uncertainty, or features of two separate stimuli) are required to distinguish different confidence models^52^. As we shall see next, however, fine discrimination tasks are not able to assess how category boundaries change with uncertainty.

#### S1.2 Incentive to shift the category boundary when uncertainty changes

In the embedded category task, the category distributions overlap in such a way that the optimal category boundaries shift when the uncertainty in the measurement distributions changes (**Figure 3a,b**). Therefore, observers have an incentive to shift their category decision rules when uncertainty changes, and we as experimenters are able to assess whether they do so.

In standard SDT tasks, in contrast, the optimal category boundary does not depend on uncertainty *σ* if the means of the internal measurement distributions remain symmetric about the boundary (**Figure S1b**). So if attention does not change the means, or changes them symmetrically (as in a discrimination task), then the optimal category boundary will not change. Observers therefore have no incentive to change their category decision rules when uncertainty changes, making it impossible to test whether the category boundary is fixed or flexible.

In summary, the embedded category task has two critical advantages over standard SDT tasks, which allow an unambiguous determination of whether and how perceptual decisions take uncertainty into account.

### S2 Modeling

#### S2.1 Lapse rates

In category and confidence models, we fit three different types of lapse rate. On each trial, there is some fitted probability of:

- A “full lapse” in which the category report is random, and confidence report is chosen from a distribution over the four levels defined by *λ*_1_, the probability of a “very low confidence” response, and *λ*_4_, the probability of a “very high confidence” response, with linear interpolation for the two intermediate levels.
- A “confidence lapse” *λ*_confidence_ in which the category report is chosen normally, but the confidence report is chosen from a uniform distribution over the four levels.
- A “repeat lapse” *λ*_repeat_ in which the category and confidence response is simply repeated from the previous trial.

In category choice models, we fit a standard category lapse rate *λ*, as well the above “repeat lapse” *λ*_repeat_.

#### S2.2 Parameterization

All parameters that defined the width of a distribution (*σ*_valid_, *σ*_neutral_, *σ*_invalid_, *σ*d) were sampled in log-space and exponentiated during the computation of the log likelihood. See **Table S1** for a complete list of model parameters for category choice and confidence models and **Table S3** for choice-only models.

#### S2.3 Visualization of model fits

Model fits were plotted by bootstrapping synthetic group datasets with the following procedure: For each model and observer, we generated 20 synthetic datasets, each using a different set of parameters sampled, without replacement, from the posterior distribution of parameters. Each synthetic dataset was generated using the same stimuli as the ones presented to the real observer. We randomly selected a number of synthetic datasets equal to the number of observers to create a synthetic group dataset. For each synthetic group dataset, we computed the mean response per orientation bin. We then repeated this 1,000 times and computed the mean and standard deviation of the mean output per bin across all 1,000 synthetic group datasets, which we then plotted as the shaded regions. Therefore, shaded regions represent the mean ±1 SEM of synthetic group datasets.

For plots with stimulus orientation on the horizontal axis (**Figures 2b**, **3c**, **S3b**, **S5a**), orientation was binned according to quantiles of the stimulus distributions so that each point consisted of roughly the same number of trials. We took the overall stimulus distribution 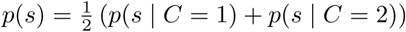 and found bin edges such that the probability mass of *p*(*s*) was the same in each bin. We then plotted the binned data with linear spacing on the horizontal axis.

#### S2.4 Model comparison metric analysis

We determined that our results were not dependent on our choice of model comparison metric. We computed AIC, BIC, AICc, WAIC^53^, and LOO for all models in the 2 model groupings (category choice-plus-confidence and category choice-only), multiplying the non-LOO metrics by 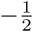 to match the scale of LOO. For AIC, BIC, and AICc, we selected the MCMC sample with the highest log likelihood as our maximum-likelihood parameter estimate. Then we computed Spearman’s rank correlation coefficient for every possible pairwise comparison of model comparison metrics for all model and dataset combinations, producing 20 total values (2 model groupings × 10 possible pairwise comparisons of model comparison metrics). All values were greater than 0.998, indicating that, had we used an information criterion instead of LOO, we would not have changed our conclusions. Furthermore, there are no model groupings in which the identities of the lowest-and highest-ranked models are dependent on the choice of metric. The agreement of these metrics strengthens our confidence in our conclusions.

#### S2.5 Model recovery analysis

We performed a model recovery analysis^54^ to test our ability to distinguish our choice and confidence models. We generated synthetic datasets from each model, using the same sets of stimuli that were originally randomly generated for each of the 12 observers. To ensure that the statistics of the generated responses were similar to those of the observers, we generated responses to these stimuli from 8 of the randomly chosen parameter estimates obtained via MCMC sampling (as described in **Section 2.4**) for each observer and model. In total, we generated 384 datasets (4 generating models × 12 observers × 8 datasets). We then fit all four models to every dataset, using maximum likelihood estimation (MLE) of parameters by an interior-point constrained optimization (MATLAB’s *fmincon*), and computed AIC scores from the resulting fits. For reasons of computational tractability, we used AIC instead of LOO as the model comparison metric. Because AIC and LOO scores gave us near-identical model rankings for data from real subjects (**Section 2.5.1**), we do not believe that the model recovery results are dependent on choice of metric.

We found that the true generating model was the best-fitting model, on average, in all cases (**Figure S4**). Overall, AIC “selected” the correct model (i.e., AIC scores were lowest for the model that generated the data) for 87.5% of the datasets, indicating that our models are distinguishable.

## Supplementary Figures

**Figure S2:**
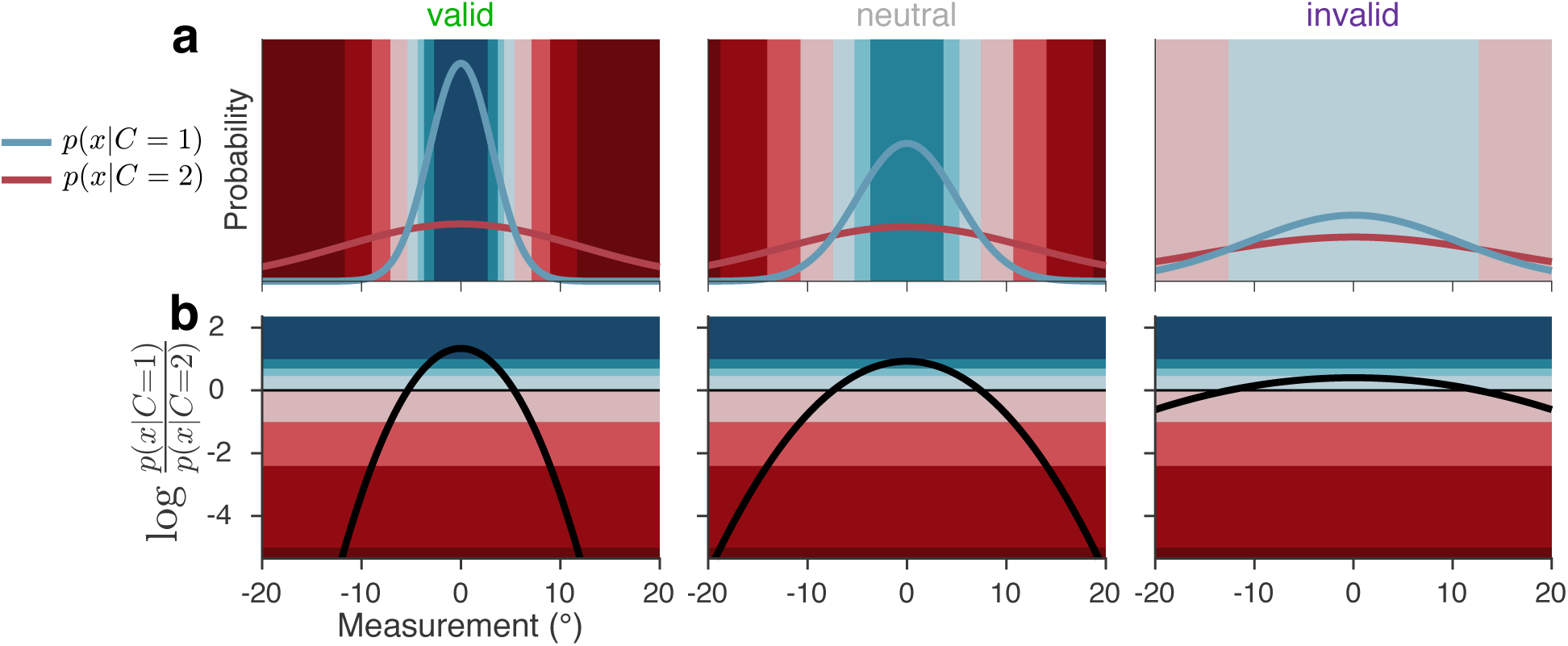
The Bayesian mapping from orientation measurement and attention-dependent uncertainty to response. Colors correspond to category and confidence response as in **Figure 1b**. (**a**) Blue and red curves show likelihood functions for the category distributions under example levels of uncertainty. (**b**) The Bayesian model maps measurement and uncertainty onto the decision variable, the log likelihood ratio (black curve). When the relative likelihood of category 1 is high, the decision variable is large and positive; when the relative likelihood of category 2 is high, it is large and negative. Response is determined by comparing the decision variable to boundaries that are fixed in log-likelihood-ratio space, but in measurement space vary as a function of uncertainty.

**Figure S3:**
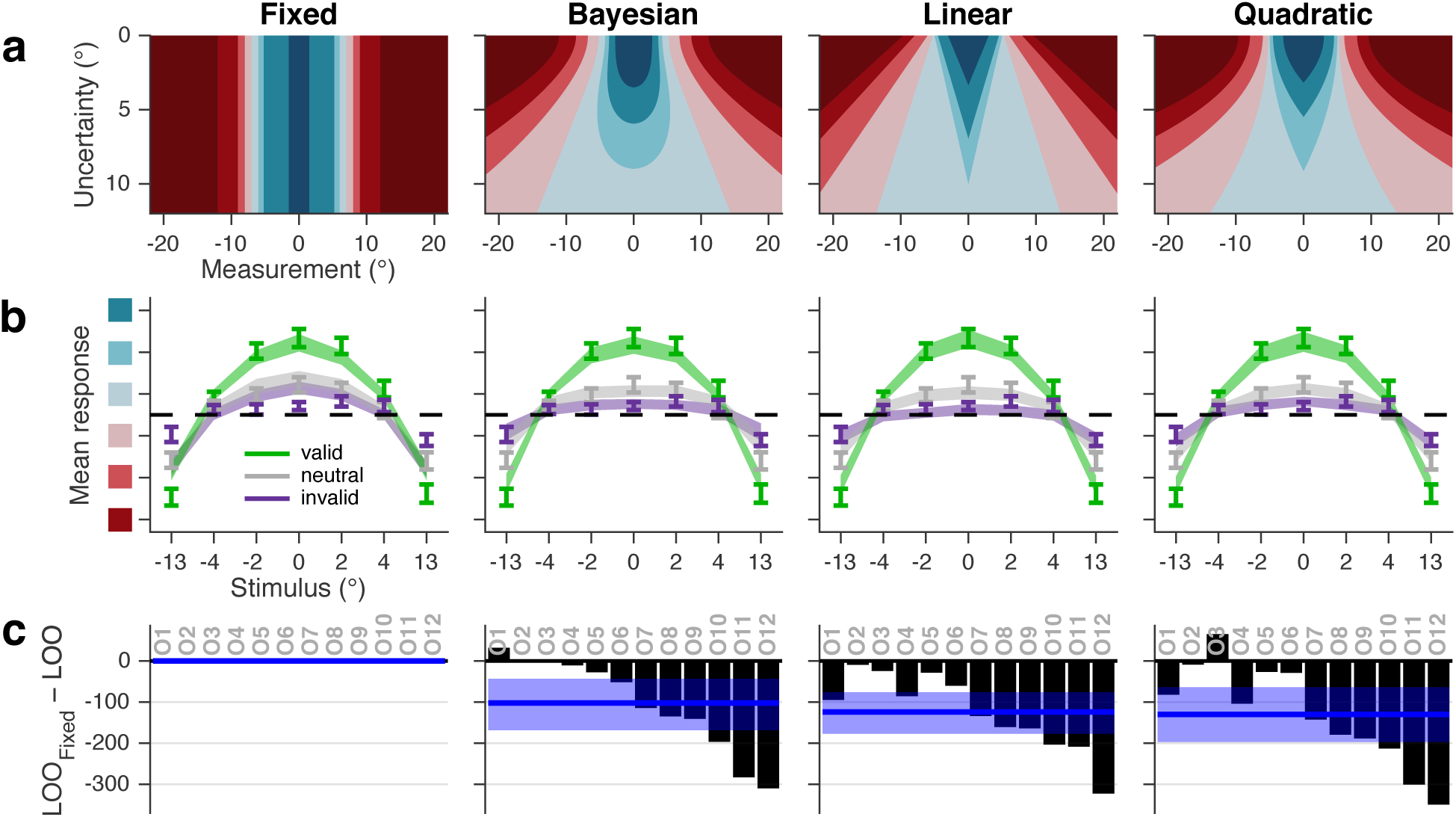
Category and confidence models. (**a**) Theoretical relation between orientation uncertainty and category and confidence decision boundaries for all models. (**b**) Mean response as a function of orientation and cue validity, as in **Figure 3c**. Stimulus orientation is binned to approximately equate the number of trials per bin. (**c**) Model comparison. Black bars represent individual observer LOO score differences of each model from Fixed. Negative values indicate that the corresponding model had a higher (better) LOO score than Fixed. Blue line and shaded region show median and 95% confidence interval of bootstrapped mean LOO differences across observers.

**Figure S4:**
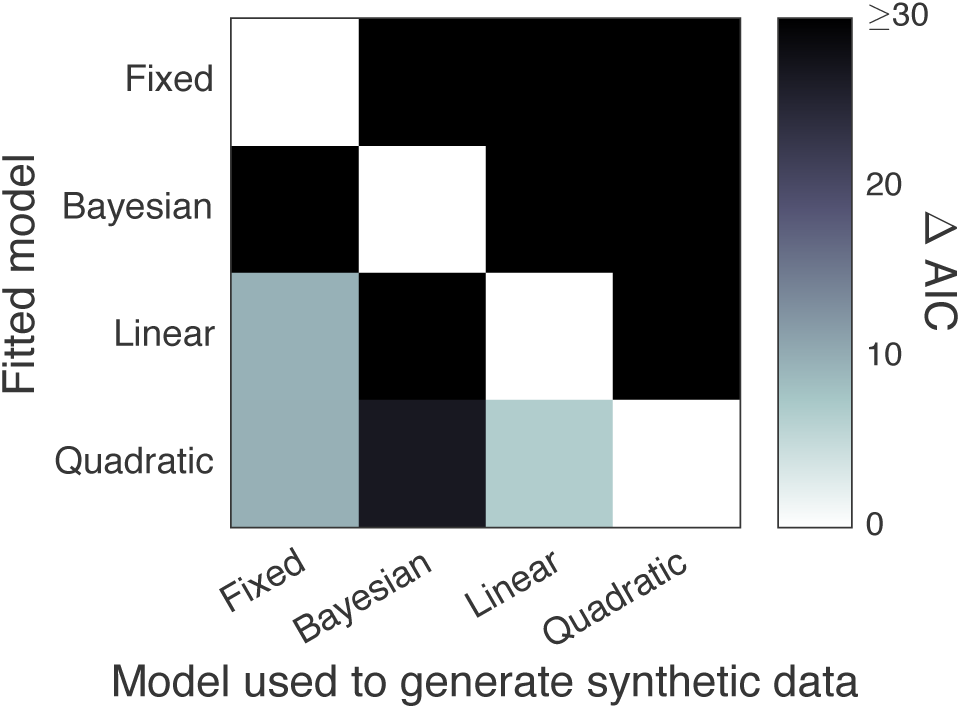
Model recovery analysis. Shade represents the difference between the mean AIC score (across synthetic datasets) for each fitted model and for the one with the lowest mean AIC score. White squares indicate the model that had the lowest mean AIC score when fitted to data generated from each model. The fact that all white squares lie on the diagonal indicates that the true generating model was the best-fitting model, on average, in all cases.

**Figure S5:**
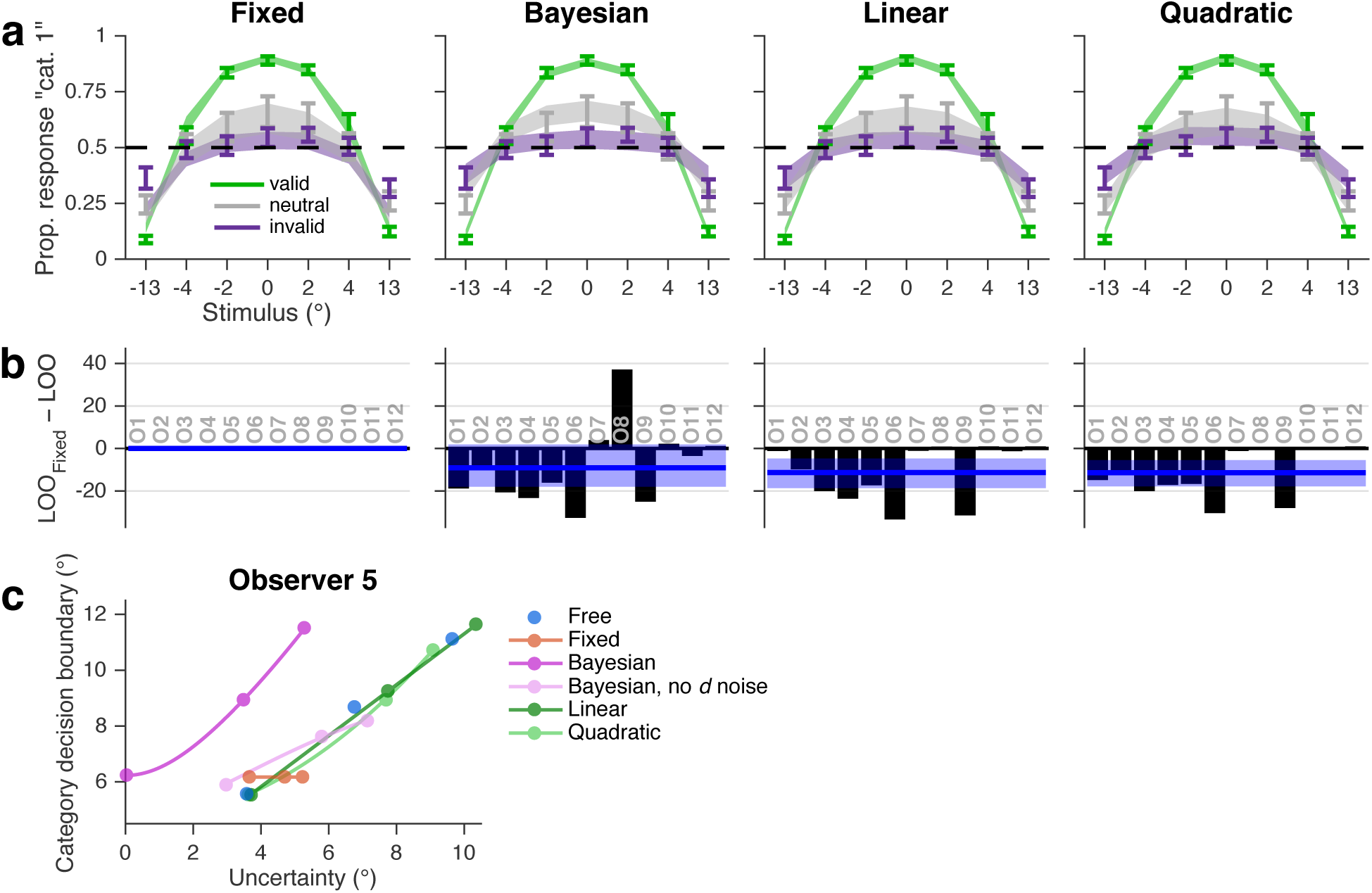
Category choice-only models. (**a**) Proportion of category 1 responses as a function of orientation and cue validity. Error bars show mean and SEM across observers. Shaded regions are mean and SEM of model fits (**Methods**). Stimulus orientation is binned to approximately equate the number of trials per bin. (**b**) LOO model comparison, as in **Figure S3c**. (**c**) Mean MCMC orientation uncertainty and category choice boundary parameter estimates for a representative observer. Estimates are plotted as a function of attention condition (valid, neutral, invalid; filled circles), along with their generating functions (curves), for the four main models fit to the category choice data only, plus a Bayesian model with no noise on the decision variable *d* and a nonparametric model in which choice boundaries are unconstrained (Free; parameter estimates from this model are plotted in gray for all subjects in **Figure 4**). The Bayesian curve is to the left of the other curves, because noise attributed to orientation uncertainty in the other models is partially attributed to decision noise in the Bayesian model; when the decision noise parameter is removed (Bayesian, no *d* noise), the curve aligns with the others.

**Figure S6:**
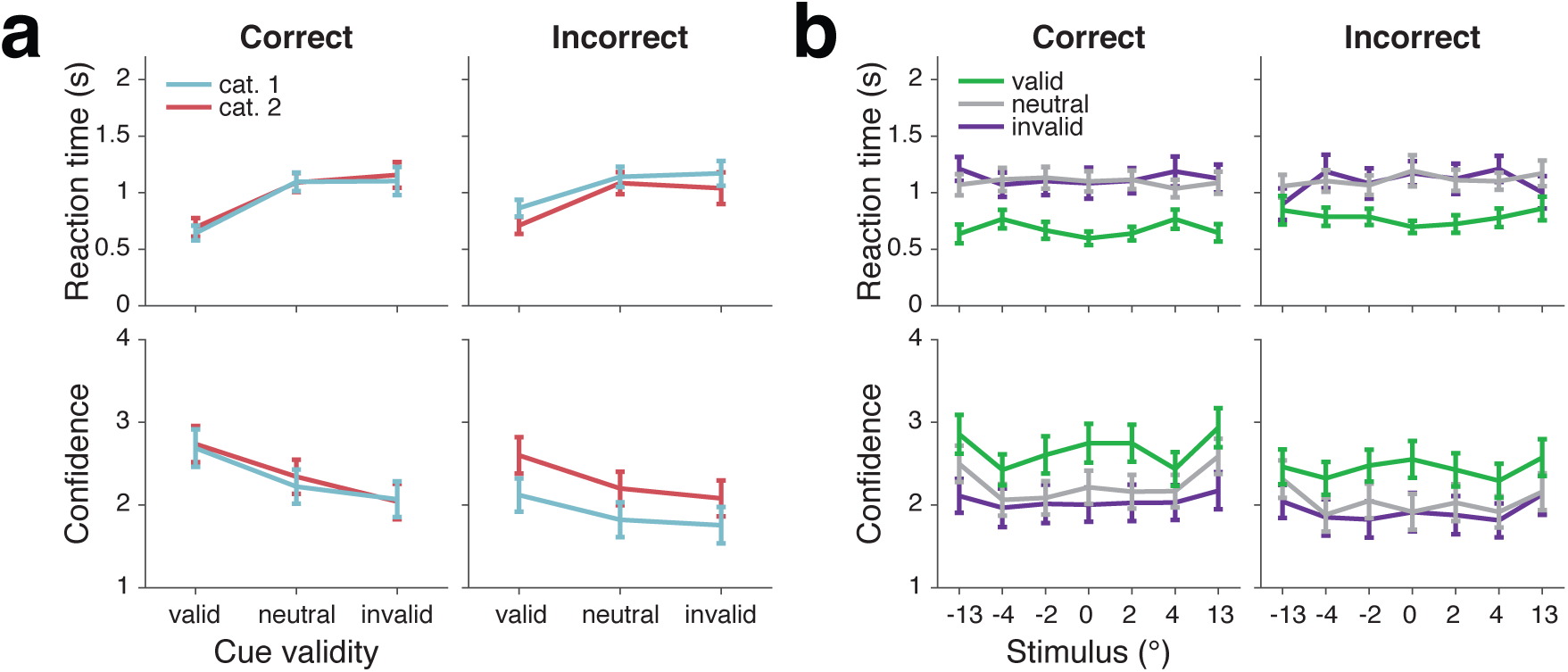
RT and confidence data broken down by category and accuracy. RT did not depend strongly on category or accuracy, though it was slightly longer for valid incorrect compared to valid correct trials. Confidence was higher overall for correct compared to incorrect trials. Confidence was higher and RT slightly faster for category 2 incorrect trials compared to category 1 incorrect trials, likely because there are more category 2 trials with high probability of being category 1 (which would lead to a high confidence error) than category 1 trials with high probability of being category 2.

**Figure S7:**
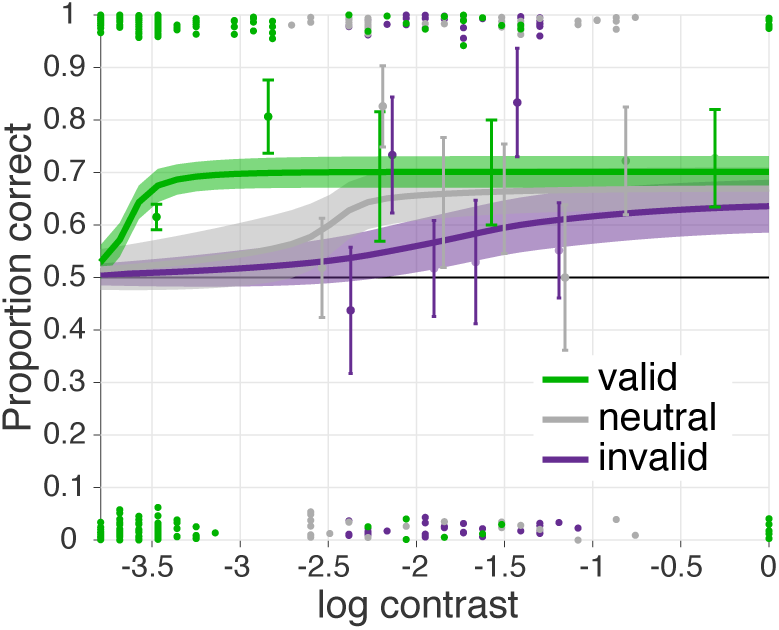
Example plot used to determine per-observer stimulus contrast. Each curve shows the mean ±1 SD of the posterior over psychometric functions for each attention condition. Error bars indicate the mean ±1 SD of the beta distribution over correctness within log contrast bins. A dot indicates one correct or incorrect trial, located respectively at the top or bottom of the plot, with vertical jitter. For this example observer, we selected a natural log contrast of −2.3 (i.e., a contrast of 10%).

## Supplementary Tables

**Table S1:** Parameters of category choice and confidence decision models.

|  | Fixed | Bayesian | Linear | Quadratic |
| --- | --- | --- | --- | --- |
| Measurement noise | $\sigma_{\text{valid}}, \sigma_{\text{neutral}}, \sigma_{\text{invalid}}$ | | | |
| Orientation-dependent noise | $\psi$ | | | |
| Decision boundaries | $k_{1-7}$ | | $k_{1-7}, m_{1-7}$ | |
| $d$ noise | | $\sigma_d$ | | |
| Lapse rates | $\lambda_1, \lambda_4, \lambda_{\text{confidence}}, \lambda_{\text{repeat}}$ | | | |
| Total number of parameters | 15 | 16 | 22 | 22 |

**Table S2:** Cross comparison of all category choice and confidence decision models. Cells indicate medians and 95% CI of bootstrapped mean LOO score differences. A positive median indicates that the model in the corresponding row had a higher score (better fit) than the model in the corresponding column.

|  |  | 15 pars.<br>Fixed | 16 pars.<br>Bayesian | 22 pars.<br>Linear |
| --- | --- | --- | --- | --- |
| 22 pars. | Quadratic | 129 [65, 198] | 27 [0, 53] | 5 [-18, 28] |
| 22 pars. | Linear | 124 [77, 177] | 21 [-3, 48] |  |
| 16 pars. | Bayesian | 102 [45, 167] |  |  |

**Table S3:** Parameters of category choice-only decision models. * indicates models that were used only for obtaining parameter estimates (**Figures 4**, **S5c**), and not for model comparison.

| | Fixed | Bayesian | Bayesian,<br>no $d$ noise* | Linear | Quadratic | Free* |
| --- | --- | --- | --- | --- | --- | --- |
| Measurement noise | $\sigma_{\text{valid}}, \sigma_{\text{neutral}}, \sigma_{\text{invalid}}$ | | | | | |
| Orientation-dependent noise | $\psi$ | | | | | |
| Decision boundaries | $k$ | | | $k, m$ | | $k_{\text{valid}}, k_{\text{neutral}}, k_{\text{invalid}}$ |
| $d$ noise | | $\sigma_d$ | | | | |
| Lapse rates | $\lambda, \lambda_{\text{repeat}}$ | | | | | |
| Total number of parameters | 7 | 8 | 7 | 8 | 8 | 9 |

**Table S4:** Cross comparison of all category choice-only decision models. Conventions as in **Table S2**.

|  |  | 7 pars.<br>Fixed | 8 pars.<br>Bayesian | 8 pars.<br>Linear |
| --- | --- | --- | --- | --- |
| 8 pars. | Quadratic | 11 [5, 18] | 2 [-2, 9] | 0 [-2, 3] |
| 8 pars. | Linear | 11 [4, 19] | 2 [-3, 10] |  |
| 8 pars. | Bayesian | 9 [-2, 18] |  |  |

## References

[1] Knill, David C & Richards, W. Perception as Bayesian Inference (Cambridge University Press, 1996).

[2] Trommershäuser, J., Kording, K. & Landy, M. S. (eds.) Sensory Cue Integration (Oxford University Press, 2011).

[3] Ma, W. J. & Jazayeri, M. Neural coding of uncertainty and probability. Annual Review of Neuroscience (2014).

[4] Qamar, A. T. et al. Trial-to-trial, uncertainty-based adjustment of decision boundaries in visual categorization. Proceedings of the National Academy of Sciences (2013).

[5] Adler, W. T. & Ma, W. J. Human confidence reports account for sensory uncertainty but in a non-Bayesian way. *bioRxiv* (2017). related:qMQe6XVts8AJ.

[6] Mamassian, P. Visual Confidence. Annual Review of Vision Science (2016).

[7] Fleming, S. M. & Daw, N. D. Self-evaluation of decision-making: A general Bayesian framework for metacognitive computation. Psychological Review (2017).

[8] Carrasco, M. Visual attention: the past 25 years. Vision Research (2011).

[9] Reynolds, J. H. & Chelazzi, L. Attentional modulation of visual processing. Annual Review of Neuroscience (2004).

[10] Carrasco, M., Penpeci-Talgar, C. & Eckstein, M. Spatial covert attention increases contrast sensitivity across the CSF: support for signal enhancement. Vision Research (2000).

[11] Lu, Z. L., & Dosher, B. A. External noise distinguishes attention mechanisms. Vision Research (1998).

[12] Anton-Erxleben, K. & Carrasco, M. Attentional enhancement of spatial resolution: linking behavioural and neurophysiological evidence. Nature Reviews Neuroscience (2013).

[13] Rahnev, D. et al. Attention induces conservative subjective biases in visual perception. Nature Neuroscience (2011).

[14] Rahnev, D. A., Bahdo, L., de Lange, F. P. & Lau, H. Prestimulus hemodynamic activity in dorsal attention network is negatively associated with decision confidence in visual perception. Journal of Neurophysiology (2012).

[15] Morales, J. et al. Low attention impairs optimal incorporation of prior knowledge in perceptual decisions. Attention, Perception, & Psychophysics (2015).

[16] Navajas, J., Bahrami, B. & Latham, P. E. Post-decisional accounts of biases in confidence. Current Opinion in Behavioral Sciences (2016).

[17] Cameron, E. L., Tai, J. C. & Carrasco, M. Covert attention affects the psychometric function of contrast sensitivity. Vision Research (2002).

[18] Rahnev, D. A., Maniscalco, B., Luber, B., Lau, H. & Lisanby, S. H. Direct injection of noise to the visual cortex decreases accuracy but increases decision confidence. Journal of Neurophysiology (2012).

[19] Caetta, F. & Gorea, A. Upshifted decision criteria in attentional blink and repetition blindness. Visual Cognition (2010).

[20] Gorea, A., Caetta, F. & Sagi, D. Criteria interactions across visual attributes. Vision Research (2005).

[21] Zak, I., Katkov, M., Gorea, A. & Sagi, D. Decision criteria in dual discrimination tasks estimated using external-noise methods. Attention, Perception, & Psychophysics (2012).

[22] Gorea, A. & Sagi, D. Failure to handle more than one internal representation in visual detection tasks. Proceedings of the National Academy of Sciences of the United States of America (2000).

[23] Gorea, A. & Sagi, D. Disentangling signal from noise in visual contrast discrimination. Nature Neuroscience (2001).

[24] Gorea, A. & Sagi, D. Natural extinction: A criterion shift phenomenon. Visual Cognition (2002).

[25] Giordano, A. M., McElree, B. & Carrasco, M. On the automaticity and flexibility of covert attention: a speed-accuracy trade-off analysis. Journal of Vision (2009).

[26] Vehtari, A., Gelman, A. & Gabry, J. Efficient implementation of leave-one-out cross-validation and WAIC for evaluating fitted Bayesian models. arXiv.org (2015). 3706900867636205788related: 3BgL-bKOcTMJ.

[27] Zizlsperger, L., Sauvigny, T. & Haarmeier, T. Selective attention increases choice certainty in human decision making. PLoS ONE (2012).

[28] Zizlsperger, L., Sauvigny, T., Händel, B. & Haarmeier, T. Cortical representations of confidence in a visual perceptual decision. Nature communications (2014).

[29] Wilimzig, C., Tsuchiya, N., Fahle, M., Einhäuser, W. & Koch, C. Spatial attention increases performance but not subjective confidence in a discrimination task. Journal of Vision (2008).

[30] Kurtz, P., Shapcott, K. A., Kaiser, J., Schmiedt, J. T. & Schmid, M. C. The Influence of Endogenous and Exogenous Spatial Attention on Decision Confidence. Scientific reports (2017).

[31] Kontsevich, L. L., Chen, C.-C., Verghese, P. & Tyler, C. W. The unique criterion constraint: a false alarm? Nature Neuroscience (2002).

[32] Dosher, B. A., & Lu, Z. L. Noise exclusion in spatial attention. (2000).

[33] Zylberberg, A., Roelfsema, P. R. & Sigman, M. Variance misperception explains illusions of confidence in simple perceptual decisions. Consciousness and Cognition (2014).

[34] Thomas, J. P. & Gille, J. Bandwidths of orientation channels in human vision. JOSA (1979).

[35] Rausch, M. & Zehetleitner, M. Visibility is not equivalent to confidence in a low contrast orientation discrimination task. Frontiers in psychology (2016).

[36] Baldassi, S., Megna, N. & Burr, D. C. Visual clutter causes high-magnitude errors. PLoS Biology (2006).

[37] Schoenherr, J. R., Leth-Steensen, C. & Petrusic, W. M. Selective attention and subjective confidence calibration. Attention, Perception, & Psychophysics (2010).

[38] Ling, S. & Carrasco, M. Sustained and transient covert attention enhance the signal via different contrast response functions. Vision Research (2006).

[39] Carrasco, M., Ling, S. & Read, S. Attention alters appearance. Nature Neuroscience (2004).

[40] Fetsch, C. R., Kiani, R., Newsome, W. T. & Shadlen, M. N. Effects of Cortical Microstimulation on Confidence in a Perceptual Decision. Neuron (2014).

[41] Peters, M. A. K. et al. Transcranial magnetic stimulation to visual cortex induces suboptimal introspection. Cortex (2017).

[42] Pelli, D. G. The VideoToolbox software for visual psychophysics: transforming numbers into movies. Spatial vision (1997).

[43] Brainard, D. H. The Psychophysics Toolbox. Spatial vision (1997).

[44] Kleiner, M., Brainard, D. H. & Pelli, D. G. What’s new in Psychtoolbox-3? ECVP Abstract Supplement. Perception (2007).

[45] Kontsevich, L. L. & Tyler, C. W. Bayesian adaptive estimation of psychometric slope and threshold. Vision Research (1999).

[46] Prins, N. The psychometric function: the lapse rate revisited. Journal of Vision (2012).

[47] Girshick, A. R., Landy, M. S. & Simoncelli, E. P. Cardinal rules: visual orientation perception reflects knowledge of environmental statistics. Nature Neuroscience (2011).

[48] Acerbi, L., Vijayakumar, S. & Wolpert, D. M. On the origins of suboptimality in human probabilistic inference. PLoS computational biology (2014).

[49] Neal, R. M. Slice sampling. Annals of statistics (2003).

[50] Acerbi, L., Dokka, K., Angelaki, D. E. & Ma, W. J. Bayesian comparison of explicit and implicit causal inference strategies in multisensory heading perception. bioRxiv (2017).

[51] Green, D. M., & Swets, J. A. Signal detection theory and psychophysics (Wiley, 1966).

[52] Aitchison, L., Bang, D., Bahrami, B. & Latham, P. E. Doubly Bayesian Analysis of Confidence in Perceptual Decision-Making. PLoS computational biology (2015).

[53] Gelman, A., Hwang, J. & Vehtari, A. Understanding predictive information criteria for Bayesian models. Statistics and Computing (2014).

[54] van den Berg, R., Awh, E. & Ma, W. J. Factorial comparison of working memory models. Psychological Review (2014).

